# Themisto: a scalable colored *k*-mer index for sensitive pseudoalignment against hundreds of thousands of bacterial genomes

**DOI:** 10.1101/2023.02.24.529942

**Authors:** Jarno N. Alanko, Jaakko Vuohtoniemi, Tommi Mäklin, Simon J. Puglisi

## Abstract

**Motivation:** Huge data sets containing whole-genome sequences of bacterial strains are now commonplace and represent a rich and important resource for modern genomic epidemiology and metagenomics. In order to efficiently make use of these data sets, efficient indexing data structures — that are both scalable and provide rapid query throughput — are paramount.

**Results:** Here, we present Themisto, a scalable colored *k*-mer index designed for large collections of microbial reference genomes, that works for both short and long read data. Themisto indexes 179 thousand *Salmonella enterica* genomes in 9 hours. The resulting index takes 142 gigabytes. In comparison, the best competing tools Metagraph and Bifrost were only able to index 11 thousand genomes in the same time. In pseudoalignment, these other tools were either an order of magnitude slower than Themisto, or used an order of magnitude more memory. Themisto also offers superior pseudoalignment quality, achieving a higher recall than previous methods on Nanopore read sets.

**Availability and implementation:** Themisto is available and documented as a C++ package at https://github.com/algbio/themisto available under the GPLv2 license.

**Supplementary information:** Supplementary data are available at *Bioinformatics* online.

## 1 Introduction

Pseudoalignment is an approximate form of sequence alignment that reports only whether a read matches to a reference sequence or not, without necessarily returning the genomic coordinates of the match. Computationally, pseudoalignment is much cheaper than regular alignment and retains accuracy at a level that is sufficient for many downstream applications. Pseudoalignment was originally used in RNA-seq quantification (Bray *et al*., 2016) to remove a computational bottleneck in aligning reads. It was then soon realized that the same methods are also applicable to the problem of metagenomic read assignment (Schaeffer *et al*., 2017; Reppell and Novembre, 2018). This led to a need for more efficient indexing methods to scale the method to large databases of bacterial reference genomes.

Recent popular pseudoalignment tools Bifrost (Holley and Melsted, 2020) and Metagraph (Karasikov *et al*., 2020a) vastly improve on the indexing performance of the original pseudoalignment implementation by Bray *et al*. (2016) in their tool Kallisto. For queries, Metagraph and Bifrost implement a threshold method of pseudoalignment that searches an index for all *k*-mers in the query sequence (typically *k* ≈ 30) and reports a pseudoalignment to an indexed genome if at least a fixed fraction *τ* of the *k*-mers are found in that genome (typically *τ* ≈ 0.8). The threshold method is slightly different from the method in Kallisto, which reports a pseudoalignment if all *k*-mers of the query are found in the index, excepting those *k*-mers that do not appear in any indexed sequence. Without the latter exception, the Kallisto algorithm corresponds to the threshold method with *τ* = 1.

In this paper we present Themisto, a new and robust pseudoalignment tool capable of scaling to data sets well beyond the limits of the tools mentioned above. Themisto implements a pseudoalignment method that allows for both thresholding and ignoring, or including, the query *k*-mers that are not found anywhere in the index. Our approach combines the threshold method of Metagraph and Bifrost with the Kallisto algorithm, providing adjustable criteria that can be tailored for different applications. We show that our approach is ideal for pseudoaligning Nanopore long-read sequencing data, where the previous methods struggle, while simultaneously achieving rapid query times and small index size. Our implementation also provides an efficient way to construct the index making use of recent advances on colored unitig extraction algorithms (Cracco and Tomescu, 2022) and is an order of magnitude faster than Bifrost and Metagraph for reference databases containing 100,000 or more bacterial genomes. These factors enable Themisto to leverage much larger databases than previous methods, thus representing a significant methodological advance in pseudoalignment.

The index structure of Themisto is a colored de Bruijn graph split into two parts, similarly to tools like VARI (Muggli *et al*., 2017), Metagraph (Karasikov *et al*., 2020a) and Bifrost (Holley and Melsted, 2020). First, the set of distinct *k*-mers of the input database is encoded using the Spectral Burrows-Wheeler transform (SBWT, Alanko *et al*. (2022)), which is a practical variant of the Burrows-Wheeler transform -based BOSS data structure of Bowe *et al*. (2012). Second, each *k*-mer is associated with a *color set*, which encodes the identifiers of reference sequences that contain the *k*-mer. Because many *k*-mers are specific to some reference sequence(s), Themisto uses different encodings for sparse and dense color sets. The mapping from *k*-mers to color sets is sparsified by only storing the mapping for a subset of *k*-mers we call *key k-mers*, a concept unique to Themisto. No information is lost by sparsifying the mapping.

### 1.1 Summary of Contributions

We summarize the main results of this article, as embodied by the Themisto succinct pseudoalignment index, as follows.

- Themisto implements a new, more robust pseudoalignment method that offers a graceful tradeoff between sensitivity and precision, while also supporting the threshold and intersection methods currently in wide use.
- Themisto is more than 10 times faster to construct than alternative indexes, enabling it to scale to vastly larger data sets. At the same time, it make judicious use of available disk and memory.
- Themisto offers consistently faster query throughput across different workloads. Only Metagraph in its fastest configuration has comparable performance, but then needs almost two orders of magnitude more working memory.
- Themisto is smaller than all other indexes with the exception of Metagraph when using a binary-relation wavelet tree, in which case it is over 100 times slower than Themisto.

To our knowledge, this study also represents the first comparison and appraisal of the main pseudoalignment methods currently in use.

### 1.2 Roadmap

This paper is structured as follows. The next section lays down notation and recalls basic concepts used throughout. Section 3 begins with a review of related methods, before detailing our new methods for pseudoalignment and the novel data structures used to implement the supporting index efficiently. Section 4 then explores the accuracy and sensitivity of different variants of pseudoalignment implemented in Themisto and competitors. Performance benchmarks are reported on in Section 5. Conclusions, reflections and outlook are then offered in Section 6.

## 2 Notation and Basic Concepts

Throughout, we use the usual notation and tools for strings and graphs, which we reiterate here for the sake of completeness and ease of reference.

A string *S* = *S*[1]*S*[2] … *S*[*n*] is a sequence of *n* symbols drawn from a fixed alphabet Σ of size |Σ| = *σ*. We denote with *S*[*i*] the character of string *S* at position *i*, where the positions are indexed starting from 1. We denote with *S*[*i*..*j*] the substring of *S* spanning positions from *i* to *j* inclusive. We refer to a string of length *k* as a *k*-mer. The concatenation of two strings *S* and *T* is denoted with *ST*. A bit vector is string with alphabet Σ = {0, 1}. A bit vector rank query, denoted rank_*B*_(*i*), for bit vector *B* at position *i* counts the number of 1-bits in *B*[1..*i*].

A set of *k*-mers can be succinctly represented as a *de Bruijn graph*. In this paper, we use the *node-centric* definition. The node-centric de Bruijn graph of order *k* for a set of strings is a directed graph (*V, E*) with vertices *V* and edges *E* such that the vertices *V* are the set of all distinct *k*-mers that are a substring of at least one of the input strings, and there is an edge from *u* to *v* iff *u*[2..*k*] = *v*[1..*k* − 1]. An edge (*u, v*) ∈ *E* is labelled with the (*k* + 1)-mer *u*[1..*k*]*v*[*k*]. The label of a path *v*_1_, … *v*_*n*_ is defined as the concatenation *v*_1_[1..*k*]*v*_2_[*k*]*v*_3_[*k*] … *v*_*n*_[*k*]. A *unitig* is a maximal path *v*_1_, …, *v*_*n*_ such that outdegree(*v*_*i*_) = 1 for all *i* = 1, …, *n* − 1 and indegree(*v*_*i*_) = 1 for all *i* = 2, …, *n*.

## 3 Methods

In this section we first outline the pseudoalignment methods currently in use. We then describe the pseudoalignment methods implemented in Themisto, which include a novel hybrid approach that combines the flexibility of resolution and strictness of previous methods. We then outline the data structures used within Themisto to efficiently implement pseudoalignment, detailing how these data structures can be scalably constructed and represented compactly while still allowing rapid query throughput.

### 3.1 Related methods

The pseudoalignment method of Themisto combines ideas from the transcription quantification tool Kallisto (Bray *et al*., 2016) and colored/annotated de Bruijn graph tools Bifrost (Holley and Melsted, 2020) and Metagraph (Karasikov *et al*., 2020a). The latter two tools implement pseudoalignment by searching for all *k*-mers of the query in the index, and then returning identifiers of all targets that contain at least *τ* percent of the *k*-mers of the query. We call this the *threshold method*. Typically, *τ* is chosen to be approximately 0.8 (which is the default value used in Bifrost; Metagraph uses *τ* = 0.7 by default). While this seems to be a good heuristic, it may not be sensitive enough to distinguish between closely related genomes that differ by only a small number of mutations in the area matching the query.

Pseudoalignment as implemented in Kallisto works approximately like that of Bifrost and Metagraph with threshold *τ* = 1; that is, *all k*-mers must be found in a reference sequence in order to report a pseudoalignment, with the important exception that *k*-mers that are not found in *any* reference sequence in the index are ignored. The rationale is that such *k*-mers (i.e., those not found in the index) are likely to be sequencing errors or sequencing artifacts such as adapters or barcodes, and thus should not be allowed to have an effect on pseudoalignment results. To implement the method efficiently, Kallisto stores the *color set* of each *k*-mer in the index, where the color set of a *k*-mer *x* contains the identifiers of all reference sequences that contain *x*. By organizing the data this way, pseudoalignment reduces to computing the intersection of all non-empty color sets of the *k*-mers contained in the query sequence. We call this pseudoalignment method the *intersection method*. In fact, as a performance optimization,

Kallisto does not compute the exact intersection, but instead skips over color sets in non-branching paths of the de Bruijn graph of the index. A skip of length *𝓁* in Kallisto is considered valid if the *k*-mer at the end of the skip in the de Bruijn graph matches the *k*-mer at distance 𝓁 in the query– otherwise Kallisto does not perform the skip. However, even if the skip is considered valid, it could affect the results if the query has variation within the skipped region. Themisto, on the other hand, never performs any skipping. On an index with 100 Salmonella genomes, we observed that Kallisto reports approximately 0.03% more hits than Themisto (see Supplement, Section 3).

### 3.2 The pseudoalignment method of Themisto

Themisto implements both the threshold method and the intersection method described above, and also a novel hybrid method that combines the best features of both methods. We formalize the hybrid method as follows. Let 𝒟 = *R*_1_, …, *R*_*n*_ be a database of reference genomes. Suppose we want to pseudoalign a query sequence *Q* using threshold *τ* ∈ [0, 1]. Let *K*(*Q*) be the list of *k*-mers of *Q* that are found in at least one reference sequence in D. We report pseudoalignment of sequence *Q* to reference sequence *R*_*i*_ if and only if at least |*K*(*Q*)| · *τ* of the *k*-mers in *K*(*Q*) are found in *R*_*i*_. Figure 1 shows an example.

**Fig. 1.**
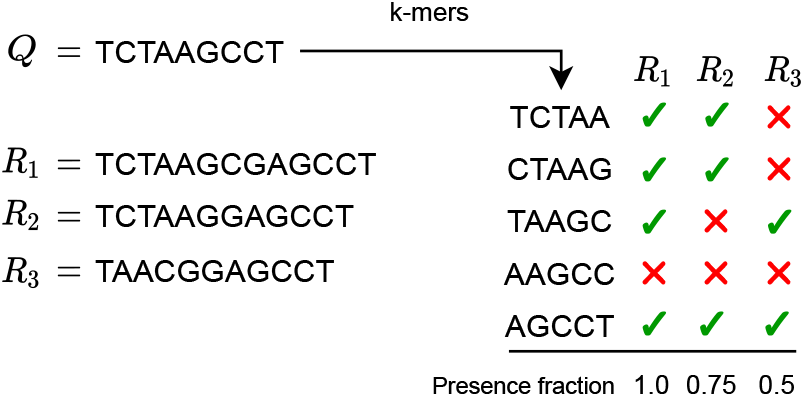
An example of the hybrid pseudoalignment method in Themisto for a query *Q* against references *R*_1_, *R*_2_ and *R*_3_ for *k* = 5. The query is first broken into *k*-mers. Then, a presence fraction is computed for each reference sequence. The presence fraction for reference *Ri* is defined as the fraction of *k*-mers of *Q* that were found in *Ri*, ignoring *k*-mers of *Q* that are not found in any reference. In this example, *k*-mer AAGCC is not present in any reference and so is ignored, making the number of *k*-mers taken into account only 4. For example, in the case of *R*_2_, three of the four relevant *k*-mers were present, so the presence fraction is 3*/*4 = 0.75. A reference is reported as found if the presence fraction exceeds the user-specified detection threshold *τ*.

The threshold *τ* can be tuned to allow a degree of variation in the pseudoalignment, while the ignoring of *k*-mers that are not in the database mitigates issues with potential sources of error — such as sequencing errors in the query, sequencing artifacts, assembly errors in the reference genomes, or an incomplete reference database — while retaining high sensitivity to variation in D. If we set *τ* = 1, the hybrid method reduces to exact computation of the intersection method of Kallisto. If we include *all k*-mers of the query, the hybrid method reduces to the threshold pseudoalignment as implemented in Bifrost and Metagraph.

### 3.3 Data structures in Themisto

The Themisto index for a given reference database 𝒟 consists of two main parts. The first part is a succinct index data structure that answers whether a query *k*-mer is found in 𝒟 and, if so, returns an integer identifier for the *k*-mer. This part of the index is implemented using the Spectral Burrow-Wheeler transform framework (Alanko *et al*., 2022), where the integer identifiers of the *k*-mers are the colexicographic ranks of the *k*-mers within the index. Note that this is a pure text index that is not aware of reverse complements, so they have to be added to the index separately. The second part is a succinct data structure that takes the identifier of the *k*-mer *x* from the first structure and uses it to retrieve the set of identifiers of the reference sequences that contain *x*. The reference sequence identifiers are called *colors* and the set of identifiers for *x* is called the *color set* of *x*. The color set of *x* is defined to also contain the colors of the reverse complement of *x*. The distinct color sets of the data are stored either as bit maps or integer arrays, depending on which one is smaller for each set.

Additional compression is achieved by only defining color sets explicitly for a specific subset of *k*-mers that we call the *key k-mers*^1^. These *k*-mers are chosen so that the color set of any other *k*-mer can be deduced from the color sets of the key *k*-mers. The idea is to select as key *k*-mers the last *k*-mer of every colored unitig, and *k*-mers around the starts and ends of reference sequences. In more detail, a *k*-mer is a key *k*-mer if the corresponding node *v* in the de Bruijn graph of the reference sequence falls into one or more of the following cases:

1. *v* has an outgoing edge into a node that is the first *k*-mer of some reference sequence.
2. *v* is the last *k*-mer of some reference sequence.
3. *v* has an outgoing edge into a node with indegree at least 2.
4. *v* has outdegree at least 2.

One of the benefits of this definition is that these conditions can be evaluated *before the color sets are constructed*, which lets us avoid constructing color sets that are not needed. Figure 2 illustrates the four cases. Let us denote de Bruijn graph nodes of the set of key *k*-mers of reference database 𝒟 with 𝒞(𝒟) and the color set of *v* with *S*(*v*). We now prove that if *v* ∉ 𝒞(𝒟), then the color set *S*(*v*) is the same as the color set of the successor of *v* in the de Bruijn graph. In the proof, the set of reference sequences also includes their reverse complements, and if a reference sequence consists of multiple disjoint sequences, each sequence is treated as a separate reference sequence in cases 1 and 2.

**Fig. 2.**
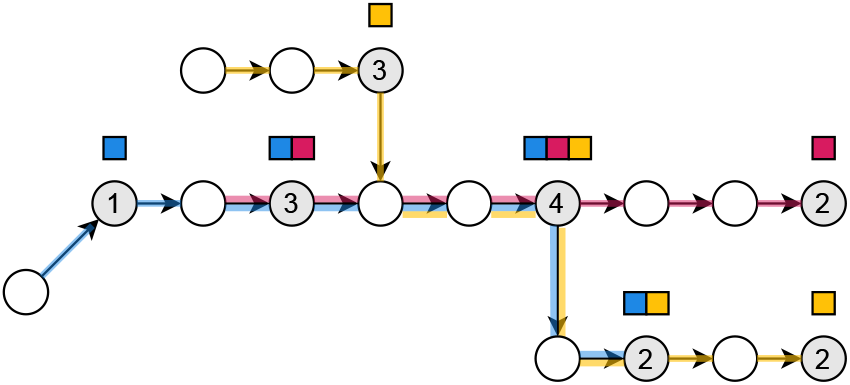
Key *k*-mers in the de Bruijn graph. The blue, red and yellow input strings are TGTTTGCTATCAC, TTGCTATCTGA and ACGTACTATCACTT respectively. The order *k* of the graph is 5. Node or edge labels are not drawn to reduce clutter. The key *k*-mers of the graph are shaded in gray, and the numbers on the gray nodes indicate which one of the four cases in the definition of key *k*-mers that node falls into. The color sets of those nodes are shown as sets of squares above the nodes. The paths taken by the sequences corresponding to the colors are shown on the edges of the graph. The color set of a node that is not shaded in gray can be obtained by walking forward to the next gray node.

Lemma 1. *If v* ∉ *𝒞(𝒟), then there is exactly one outgoing edge* (*v, u*) *and S*(*v*) = *S*(*u*).

Proof. Since case 2 does not hold for *v*, it is not the last *k*-mer of any reference sequence. This means that the outdegree of *v* is at least 1. Since case 4 does not hold for *v*, it has outdegree at most 1, so the outdegree of *v* is exactly 1. Denote the unique outgoing neighbor from *v* with *u*. Any color *c* of *S*(*v*) must also be in *S*(*u*), because otherwise the sequence of color *c* ends at *v* contradicting case 2. Symmetrically, any color *c* of *S*(*u*) must also be in *S*(*v*), since the indegree of *u* is 1 by case 3, and no new new reference sequence starts at *u* by case 1. Therefore *S*(*u*) = *S*(*v*). □

By Lemma 1, the color set of any node *u*∉𝒞(𝒟) can be obtained by walking forward to the nearest node that is in 𝒞(𝒟). By case 4, every node on such a walk has outdegree 1, so there is exactly one such walk from *u*, and by case 2, the walk is guaranteed to terminate.

The color index of Themisto stores the distinct color sets in a table, and for each node *v* ∈ 𝒞(𝒟) stores the index of the color set of *v* in the table. The rest of the color sets are inferred on demand at query time by walking forward in the graph as explained above. However this walk can be as long as the longest reference sequence in the worst case, so Themisto also stores the color set indices on every *d*-th node on the walks, for a tunable parameter *d* that trades space for time. These added sampled *k*-mers guarantee that at most *d* graph traversal operations are required to retrieve the color set of any *k*-mer.

The index is constructed by first preprocessing the input database 𝒟 into colored unitigs. We use the GGCAT tool (Cracco and Tomescu, 2022), which is a highly optimized and parallel method for colored unitig construction. This can reduce the size of the data by multiple orders of magnitude for repetitive collections of genomes. Next, we build the *k*-mer index using the SBWT construction algorithm on the set of unitigs (Alanko *et al*., 2022). Then we proceed to *mark* the key *k*-mers in a bit vector by evaluating the four conditions described above using the SBWT index. Next, we use the SBWT search routine to search for all *k*-mers of each unitig. If a searched *k*-mer has been marked as key, we store the color set in the color set table, and the mapping from the *k*-mer identifier to the color set index. Finally, we add additional *k*-mer-to-color-set mappings on every *d*-th node on non-branching paths using the SBWT to walk the paths in the graph.

The mapping from key *k*-mers and the additional sampled *k*-mers to color sets is represented by storing the color set indices only for those *k*-mers, and using bit vector rank queries to retrieve the color set index of the *i*-th such *k*-mer. As the color sets are of variable length and have different binary representations depending on the sparsity of the set, the color sets themselves are stored by concatenating their binary representations and storing in a separate array the starting position of each color set in the concatenation. Figure 3 illustrates the index structure.

**Fig. 3.**
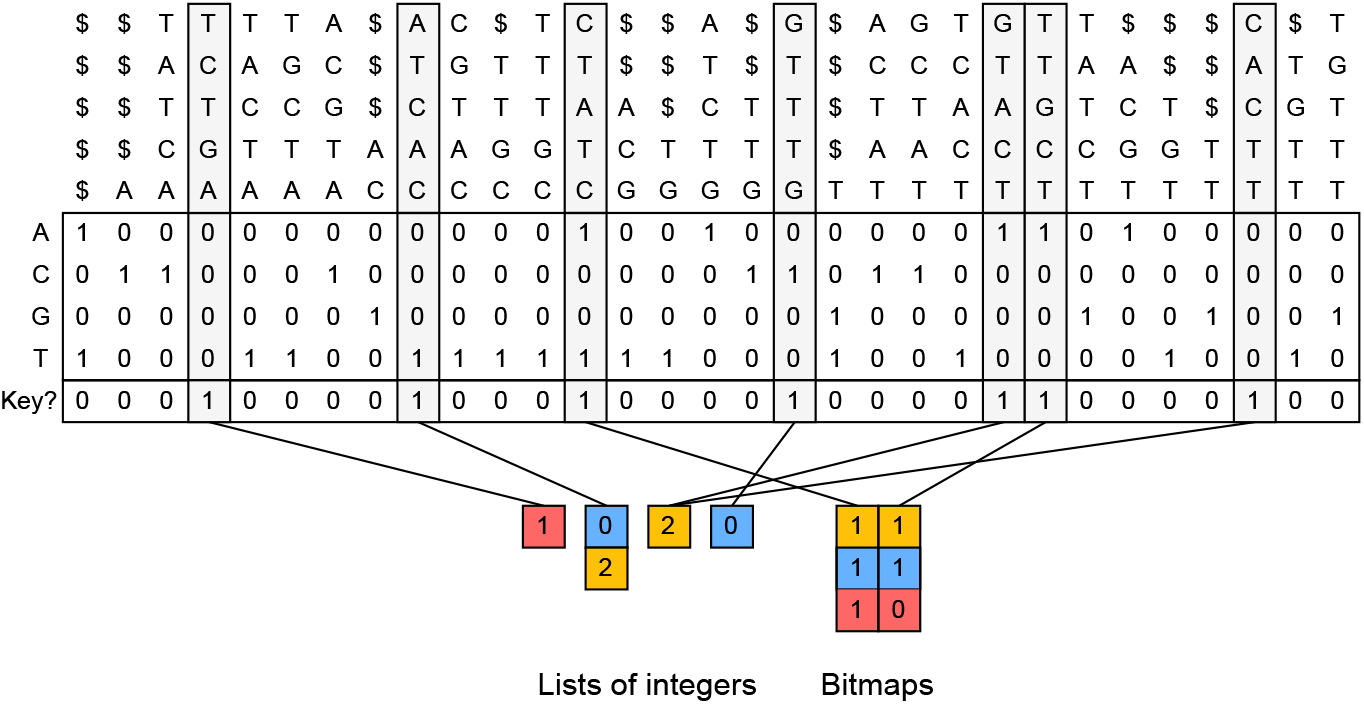
The Matrix-SBWT *k*-mer index and the mapping to color sets. This Figure continues from the example in Figure 2. The columns of the SBWT matrix represent the *k*-mers of the input data, with technical dummy prefixes containing dollar-symbols added to the *k*-mers ending in the first *k* positions of the input sequences. The *k*-mers are shown vertically at the top (for illustration purposes only – they are not excplicitly stored), and the SBWT matrix is the binary matrix in the middle with 4 rows. Each row corresponds to a character of the alphabet, and a 1-bit at cell (*i, j*) indicates that the *j*-th *k*-mer has on outgoing edge such that the last character of the edge (*k* + 1)-mer is the *i*-th character of the alphabet. See Alanko et al. (2022) for a more in-depth explanation. The columns shaded in gray are the key *k*-mers, which are also marked in the bit vector below the SBWT matrix. The key *k*-mers are associated with the color sets at the bottom. The sparse sets are encoded as lists of integers, whereas the dense sets are encoded as bit maps. The mapping from key *k*-mers to the color sets, that is represented by lines in the Figure, is implemented by marking with another bit vector (not pictured) whether the set is sparse or dense, and using a bit vector rank query to find the index of the set within the color sets of its type (sparse or dense). Color sets of a single type are stored in concatenated form, with pointers to the starts of the sets.

### 3.4 Color set compression

Themisto uses different representations for sparse and dense color sets. Sparse sets are stored simply as sequences of color identifiers, where each identifier is encoded with ⌈log *m*⌉ bits, where *m* is the number of distinct colors in the dataset. Dense sets are encoded with bit maps of length *m*, where a 1-bit at position *i* indicates the presence of color *i*, and a 0-bit indicates the absence of the color. Themisto selects the more efficient encoding for each color set separately.

The elements of sparse sets are stored sorted in ascending order to support fast set intersection in a single scan over the sorted color sequences. The dense sets can likewise be intersected very quickly by computing the bitwise AND-operation between the bit maps. The intersection between a sparse and a dense set is performed by iterating the elements of the sparse set and checking the presence of each element in the dense set with a constant-time bitmap lookup.

Themisto also includes the option to encode the color sets with Roaring bitmaps (Chambi *et al*., 2016). This representation can encode large bit maps by splitting the bit map into blocks and selecting the most effective encoding for each block. However, it encodes all bit maps with less than 4096 elements as sparse, no matter how dense the bit map is, so it is ineffective for colorings with less than 4096 colors.

## 4 Use cases

One of the main applications of Themisto is in metagenomics, where Themisto is used as part of the mGEMS pipeline (Mäklin *et al*., 2021) to identify the composition of a set of sequencing reads at the level of lineages within a bacterial species. This application is enabled by Themisto’s index structure and construction algorithm being able to scale to huge reference databases, and also by its efficient implementation of the hybrid pseudoalignment method.

The version of Themisto originally used in the mGEMS pipeline only implemented the intersection method of pseudoalignment. Our aim here is then to investigate the impact of using the threshold and hybrid methods and to guage the quality of results produced by all three methods. To this end, we produced pseudoalignments with various parameters from a single sample containing short-read (150bp) paired-end Illumina MiSeq550 reads (accession number SRR14189471). This sample contains DNA from three different lineages of *Escherichia coli* sequence type (ST) 131 in measured (i.e., known) proportions, making it ideal for a metagenomics comparison. For analysis on the effectiveness of pseudoalignment on various datasets, we point the reader to Mäklin *et al*. (2020).

We pseudoaligned the reads from the sample against a publicly available reference database (doi: 10.5281/zenodo.6656897, derived from the data published in Horesh *et al*. (2021) and Gladstone *et al*. (2021), described in Mäklin *et al*. (2022)) and ran the relative abundance estimation part of the mGEMS pipeline, called mSWEEP (we used v1.6.2 with default options, see (Mäklin *et al*., 2020)), to infer relative abundance estimates from the pseudoalignments. These were then compared to the true values.

Figure 4 shows that the performance of the intersection method (*k* = 31) was in line with using the threshold or hybrid method with a high threshold (*τ* ∈ {0.9, 0.99, 1}) when evaluated with the absolute or relative error in the estimates. Lowering the threshold (*τ* ∈ {0.7, 0.5, 0.1}) predictably reduced the accuracy of the estimates, resulting in more relative abundance assigned to the “noise” category, which contains all estimates for the lineages which were not present in the sample, and in a reduction of accuracy for all three target lineages.

**Fig. 4.**
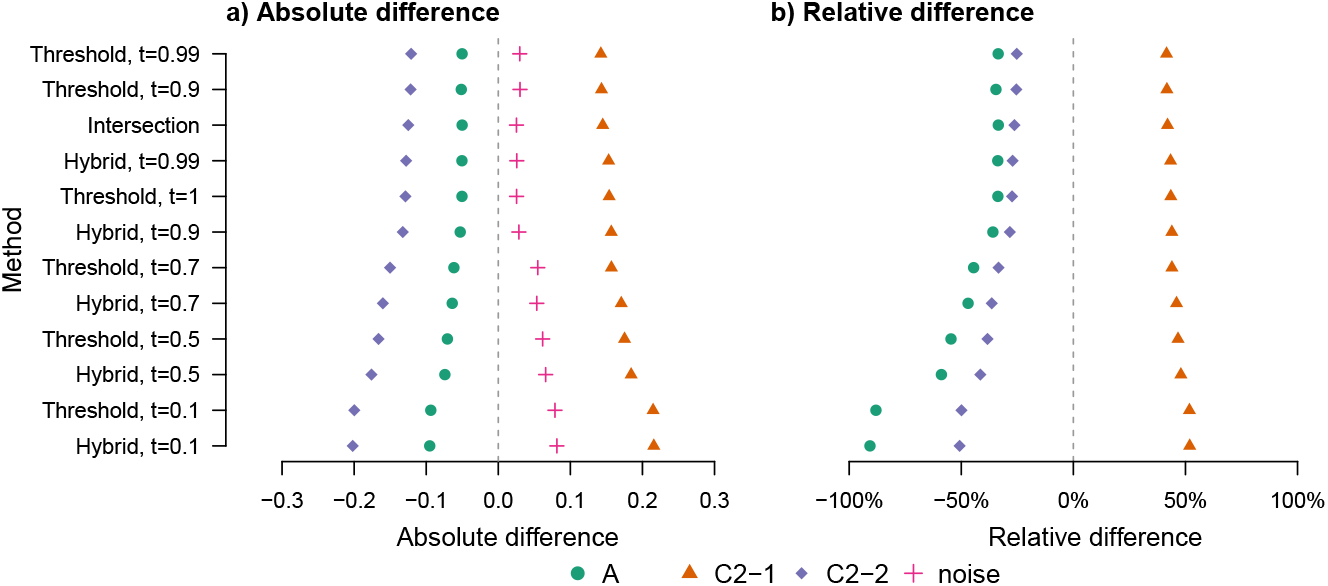
Accuracy of relative abundance estimation with different pseudoalignment criteria. The figure shows the absolute (panel a)) and relative (panel b)) difference in relative abundance estimates for three lineages of E. coli ST131. The three lineages (A, C2-1, C2-2) were sequenced together at known proportions using Illumina NextSeq550 operating on a 2×150 bp paired-end set-up. In panel a), the values labelled “noise” correspond to the sum of all estimates that are not for the three truly present lineages. In panel b), the relative difference is defined as the difference between the estimate and the true value, divided by the estimate. The vertical axis is sorted according to the sum of squared errors, with the lowest error at the top.

We next tested the different methods on pseudoaligning metagenomic long-read sequencing data with a synthetic mixture containing Nanopore sequencing reads from the same bacteria as in Figure 4. This mixture contained Nanopore reads from each of the three lineages, sequenced individually in an earlier study ((Mäklin *et al*., 2021), reads available from Zenodo with doi: 10.5281/zenodo.4738983) that were computationally mixed at the same proportions as in the short-read sequencing sample. Because the relative abundance estimation part in the mGEMS pipeline is designed for short-read data, we evaluated the accuracy of the different alignment methods based on recall of reads pseudoaligning against the 100% reference genomes. The pseudoalignment of a read was considered correct if it contained at least one pseudoalignment to any reference sequence from the same lineage and incorrect if it did not align to any sequence from the same lineage.

Pseudoalignments from the Nanopore data revealed stark differences in the recall rates of the hybrid and the threshold methods (Figure 5). The hybrid method was able to pseudoalign the reads at a high recall rate when using a threshold of 0.9 or 0.7, whereas the threshold method only performed comparably at *τ* = 0.1, which is low enough that nearly all of the reads pseudoalign to all reference sequences. Using the threshold method with a high threshold (*τ* ∈ {0.7, 0.9, 0.99, 1}) resulted in nearly zero alignments. The observed differences between the methods are likely a result of the higher error rate in Nanopore sequencing reads and the longer read length. Long reads can span multiple contigs in a reference sequence with gaps in parts of the genome that are difficult to assemble. Together, these factors result in an abundance of *k*-mers that are not present in the reference database, which makes the hybrid method essential for pseudoaligning long read data.

**Fig. 5.**
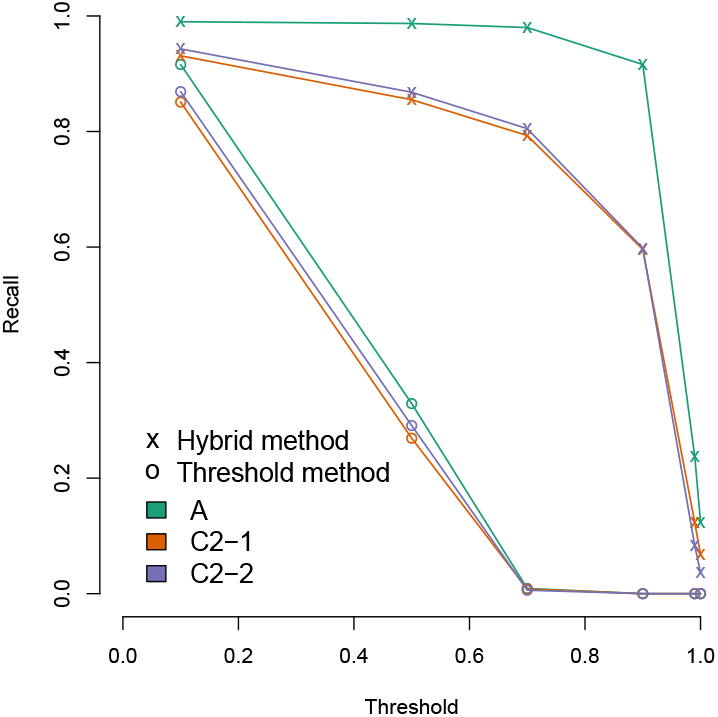
Comparison of pseudoalignment recall with the hybrid method and the threshold method at varying thresholds. The figure shows the recall in pseudoaligning a computational mixture of 100 000 Nanopore isolate sequencing reads from three lineages of E. coli ST131 at varying proportions (C2-2: 0.60, C2-1: 0.20, A: 0.20, indicated by the colors). The pseudoalignment of a single read is considered correct if it contains at least one hit against a reference genome belonging to the same lineage that the read was originally sequenced from. Recall is defined as the number of reads that align correctly divided by the total number of reads from the lineage.

For best results, we recommend using a two-step pipeline that first aligns the reads to a broad set of references colored by species, and then proceeds to pseudoalign the reads against strain-level indexes from each identified species. See Supplement, Section 2 for a discussion on advantages of this approach.

## 5 Performance

In this section, we evaluate the construction and query performance of Themisto. We compare Themisto against the two state-of-the-art tools, Bifrost (Holley and Melsted, 2020) and Metagraph (Karasikov *et al*., 2020a).

All our experiments were conducted on a machine with four 2.10 GHz Intel Xeon E7-4830 v3 CPUs with 12 cores each for a total of 48 cores, 30 MiB L3 cache, 1.5 TiB of main memory, and a 12 TiB serial ATA hard disk. The OS was Linux (Ubuntu 18.04.5 LTS) running kernel 5.4.0-58-generic. Themisto was compiled using `g++` version 10.3.0, Bifrost using g++ version 9.4.0 and for Metagraph we used the pre-compiled binaries distributed by the authors. During construction, all runtimes and RSS memory peaks were recorded using the command `/usr/bin/time` with the `-v` flag. At query time, the time to load the index into memory was disregarded and only the time spent running queries was included in the running time. The time to load the index in each tool was extracted by observing the time stamps in the log files printed by the programs themselves. In the case of Bifrost, the source code was modified to print a time stamp before and after the section of the code that runs the queries.

### 5.1 Index construction

We use a value of *k* = 31 for all experiments^2^. Both Bifrost and Metagraph split the index into two components: a *k*-mer index with support for *k*-mer lookups, and a color index that is able to return the color set of a *k*-mer after a *k*-mer lookup. The *k*-mer index of Bifrost is essentially a minimizer index (Roberts *et al*., 2004) of the set of unitig strings of the de Bruijn graph, whereas the *k*-mer index of Metagraph uses the BWT-based BOSS data structure (Bowe *et al*., 2012). For the color index, Bifrost uses bitmaps, which are compressed either using a method based on Roaring bitmaps (Chambi *et al*., 2016), or as a single 64-bit word, depending on sparsity.

Metagraph on the other hand implements a variety of different coloring structures with different time-space tradeoffs designed for different types of data and queries. We ran our experiments using the *row-major* representation, which stores a color set for each *k*-mer. This is similar to the representation in Themisto and is appropriate for pseudoalignment queries which require retrieval of color sets of individual *k*-mers. Metagraph offers variety of methods to further compress the row-major color set data structure. The user manual recommends using the `RowDiff<Multi-BRWT>` method for large instances, which is said to achieve best compression while still providing good query performance. This representation is based on the multiary binary relation wavelet tree of Karasikov *et al*. (2020b). The `RowDiff<Multi-BRWT>` representation must be constructed by constructing the column-major representation first, and transforming it to the desired form. We include in our construction benchmark both the regular row-major representation and the `RowDiff<Multi-BRWT>` representation built via the column-major representation. The reported time is the sum of running times of all construction steps, and the reported peak space is the maximum peak space of all steps. The exact command-line parameters used in the different steps can be found in the Supplement, Section 4.

Index construction performance was evaluated on increasingly large sets of *Salmonella* genomes. Our dataset was created by extracting the *n* = 178, 984 high-quality Salmonella enterica assemblies from the curated bacterial reference genome dataset of Blackwell *et al*. (2021) and has 859,337,174,122 nucleotides. We add *k*-mers of both DNA strands to the index. Since the color sets in this data set are very large, the index size will be dominated by storage of the distinct color sets. Thus, we can afford to disable the *k*-mer-to-color-set-id sparsification feature of Themisto by setting the parameter *d* described in Section 3.3 to 1. This increases the total space of the coloring structure by only 2%. In other applications, where the color sets are smaller, such as when coloring genomes by *phenotype* (Jaillard *et al*., 2018) or by species, the sparsification would have a larger impact on reducing overall space. For example, the color structure of the species-colored Themisto index of the 640k high-quality assemblies of the entire Blackwell et al. 661k genomes dataset is 90% smaller with *d* = 20 compared to *d* = 1, and queries are slightly *faster* on the smaller index. This index is available on Zenodo at https://doi.org/10.5281/zenodo.7736981. More details on this index can be found in the Supplement, Section 1.

We compared the scalability of indexing with the different methods by indexing subsets containing the first ⌊*n/*2^*i*^⌋ genomes for *i* = 0, 1, … ⌊log *n*⌋. Runs exceeding 24 hours were terminated. Figure 6 shows the indexing time, peak memory and disk-resident size of the de Bruijn graph and coloring components of all indexes. Themisto showed by far the best scalability in time, taking an order of magnitude less time than the other two tools for input sizes exceeding 10^4^ genomes, and was the only tool capable of indexing the whole dataset in under 24 hours (at which point we terminated both Bifrost and Metagraph). This high performance is in part due to the tight integration of the GGCAT tool of Cracco and Tomescu (2022) into Themisto for colored unitig extraction.

**Fig. 6.**
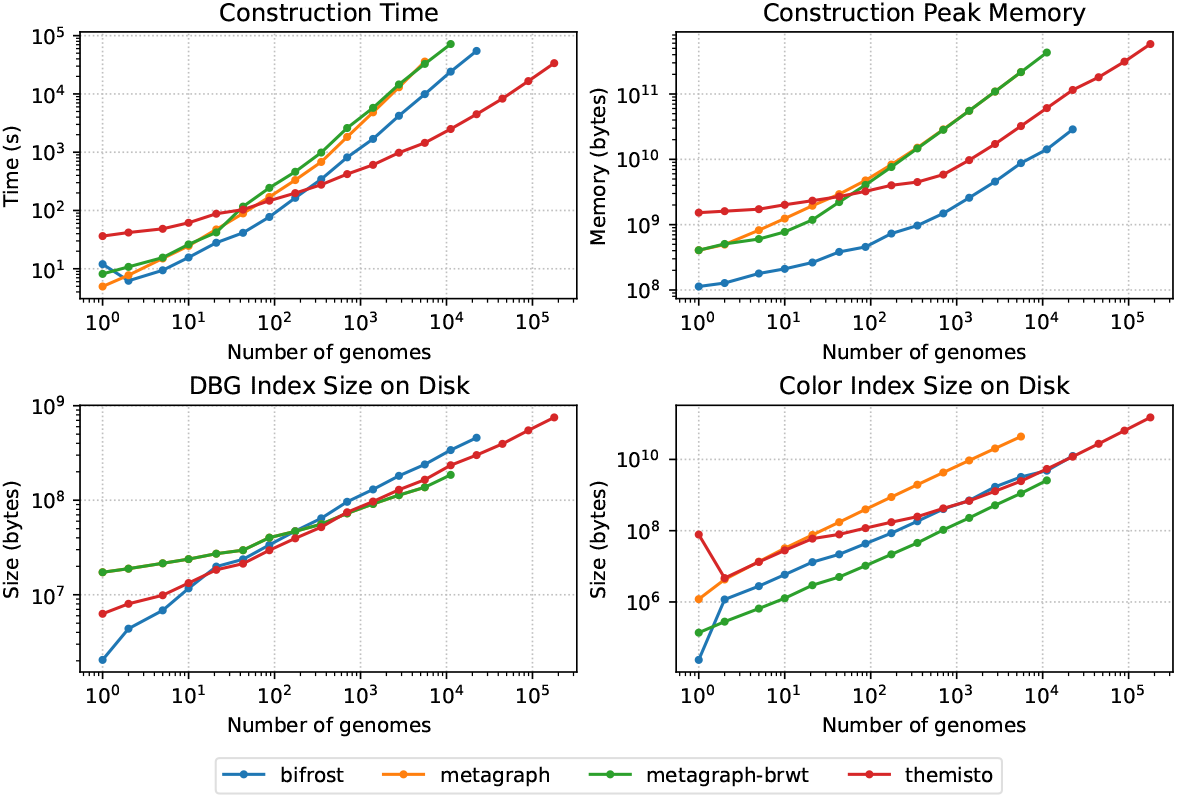
Construction time, peak memory and index size on disk. Metagraph-brwt is the binary relation wavelet tree variant of Metagraph. The DBG index is the same for both Metagraph and Metagraph-brwt, so the line for Metagraph is hidden under the line for Metagraph-brwt in the DBG index size plot (bottom left). The data in this figure is listed in Table 1.

The peak working memory of Themisto for large inputs was in between the two tools. The de Bruijn graph data structures of Themisto and Metagraph were of similar size on disk, which is to be expected as they both use a similar Burrows-Wheeler transform based index structure. The de Bruijn graph structure of Bifrost grew slightly larger than the other two on large inputs (see Figure 6, bottom-left), taking 1.3 times more space than Themisto and Metagraph. The Bifrost index on disk consists of a gzipped plain text GFA-file containing the unitigs and the edges of the graph, and a small index data structure that operates on top of the unitigs. The unitig file takes approximately 90% of the total space of the graph index without colors. This means that the *k*-mer indexes of Themisto and Metagraph take less space than a gzipped GFA description of the de Bruijn graph with no index included.

**Table 1.**
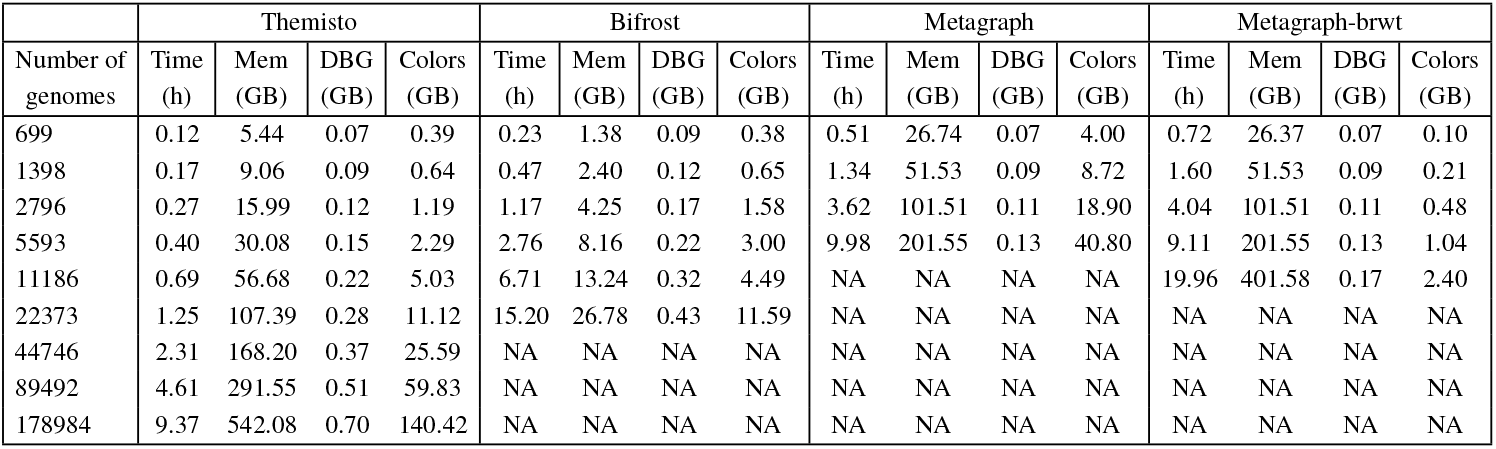
Construction running time, peak memory, and the size of the de Bruijn graph (DBG) and color components of the index structures on disk. Data for runs which exceeded the time limit of 24 hours are marked with NA. To save space, rows with less than 699 genomes have been omitted. The full table can be found in the Supplement, Section 5. The full data is also plotted in Figure 6.

The coloring structures of Themisto and Bifrost were approximately the same size for large inputs. The version of Themisto using Roaring bitmaps had a similar size, being just 3% smaller than the default Themisto color representation on the complete data set. The uncompressed row-major coloring structure of Metagraph was roughly 15 times larger than Themisto and Bifrost on large inputs, while the compressed `RowDiff<Multi-BRWT>` structure was 2.5 times smaller.

In all tools, the space requirements for storing the complete colored de Bruijn graph index were dominated by the requirements for storing the color information. With this in mind, Figure 7 shows statistics on sizes of color sets explicitly stored in the Themisto index. The subfigure on the left shows the number of color sets stored for each cardinality. We see that the most common cardinality is 2. For sets with cardinality greater than 2, the decreasing slope of the curve in log-log space stays roughly the same, indicating a power law distribution. However this trend breaks in the very largest color sets. The subplot on the right depicts the total number of bits allocated to sets of a given cardinality. Here we see that even though most of the sets in the index are very small, they are efficiently represented in the sparse encoding, and most of the space of the index is contributed by the densely encoded larger color sets. We believe that the binary relation wavelet tree in the Metagraph-brwt index is able to compress the largest color sets better than Themisto, explaining the space advantage over Themisto. However, this comes at a crippling cost to query time, as we see in the next section.

**Fig. 7.**
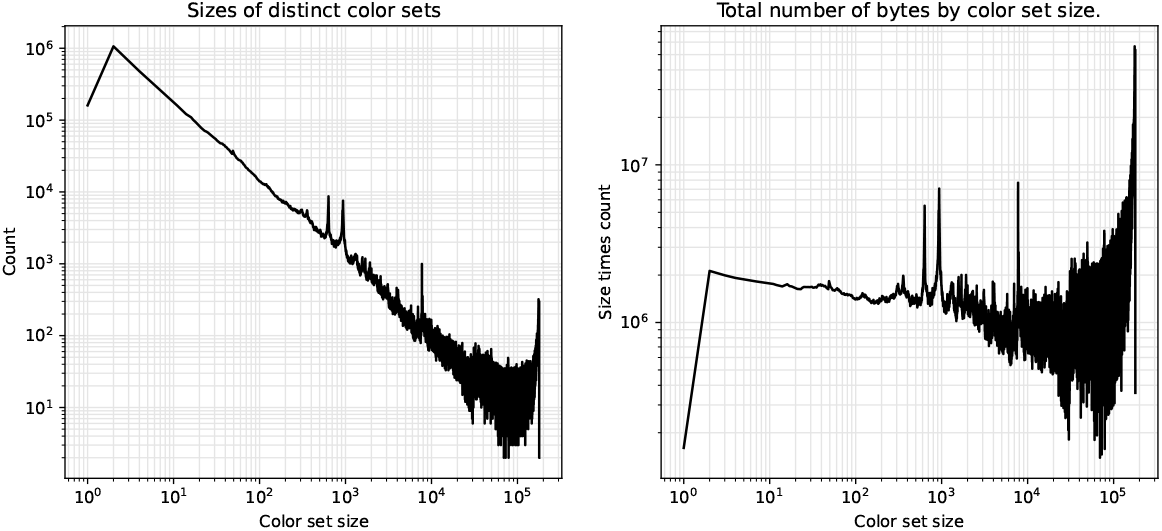
Statistics on distinct stored color sets in Themisto. The left subplot shows the number of color sets of a given cardinality. The right subplot shows the total number of bytes used for color sets of a given cardinality.

### 5.2 Queries

We first benchmarked the queries on reads sequenced from *S. enterica*. The reads were obtained from the The University of Warwick/University College Cork 10,000 *Salmonella* genomes project (Achtman *et al*., 2020). The query experiment was run using sample ERR2693452, against indexes built on the largest dataset all tools were able to process in 24 hours, which contains 5,593 genomes. The reads originate from an Illumina HiSeq X Ten machine, and have length 151, with an average FASTQ quality score of 39.

Since the reads are sampled from a *S. enterica* isolate, they accordingly match with many *Salmonella* reference genomes. This results in a very heavy workload, and the runtime is consequently dependent on the efficiency of the chosen coloring data structure. To keep the running time and the size of the output reasonable, we only align the first 500,000 reads. In all experiments in this section, the time to load the index into memory is not included in the reported times.

The hybrid pseudoalignment method of Themisto assigns 99.7% of the reads to at least one reference genome. We call reads that have at least one match “positive”. On average, a positive read was assigned to 2,275 genomes. Using the pure threshold-based pseudoalignment method *without* ignoring unknown *k*-mers results in positivity rate of only 90.0%, with 2,222 genomes reported on average per positive read. This rate is consistent with Metagraph and Bifrost, which had positivity rates of 89.7% and 90.2% respectively, reporting on average 1,111 and 2,249 genomes respectively, per positive read.

The left subplot of Figure 8 shows the query throughput and memory usage of the three tools. Themisto and Metagraph with the uncompressed row-major representation had the best throughput by an order of magnitude. However, the RAM usage of Metagraph with this representation was an order of magnitude higher than that of Themisto. Using the binary relation wavelet tree variant of Metagraph achieved the lowest memory usage in this benchmark (half the size of Themisto) but the query time was *two* orders of magnitude slower than Themisto and the uncompressed Metagraph index. Metagraph also includes a `--fast` option that can be used to speed up queries on a compressed index by uncompressing the part of the index that intersects with the query batch, but this comes at a cost of increased space usage. In this benchmark enabling the `--fast` option sped up the Metagraph queries on the binary relation wavelet tree index by a factor of 51, but increased the space usage by a factor of 23 from 1.25GB to 29GB.

**Fig. 8.**
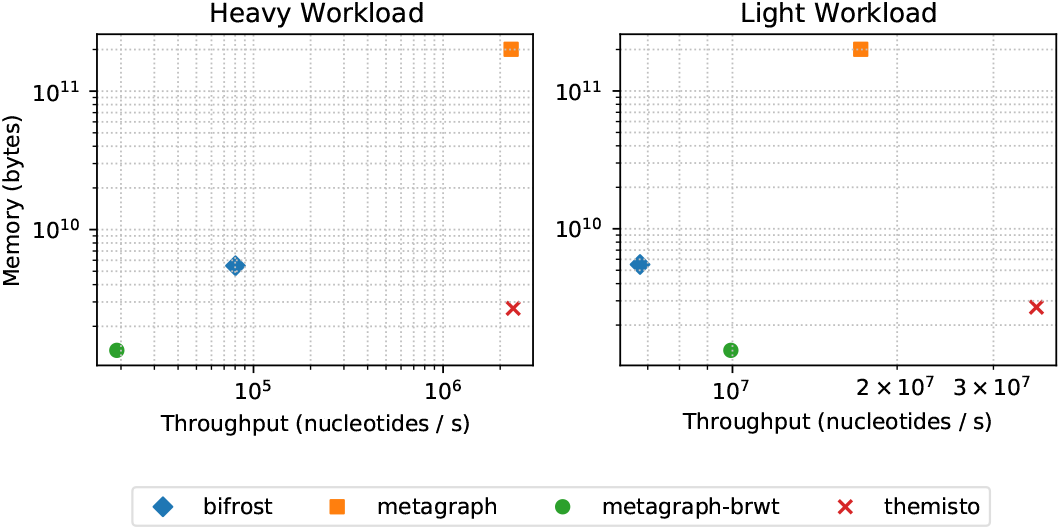
Query throughput and space. The heavy workload is the Salmonella enterica isolate query read set, and the light workload is the N. gonorrhoeae isolate query set. Metagraph-brwt is the binary relation wavelet tree variant of Metagraph. The data for these plots is listed in Table 2.

Since this workload is dominated by the processing of the color sets, we also ran the queries on reads which are unlikely to pseudoalign to *S. enterica* strains, representing a low workload case. We used reads from *Neisseria gonorrhoeae*, which is a species evolutionarily far-removed from *S. enterica*, residing in a different class in the taxonomic tree of bacteria. These reads were obtained from a whole-genome sequencing analysis study of *N. gonorrhoeae* isolates in China (Peng *et al*., 2019). The query experiment was run using sample SRR6765295 against the same *Salmonella* indexes as previously. These reads originate from an Illumina NextSeq 550 machine, and have length 101 with an average fastq quality value of 33. The positivity rate was now only 2% using the hybrid method of Themisto, and 0.6% by using the pure threshold method (Metagraph 0.6%, Bifrost 0.7%). A positive read pseudoaligned against 243 genomes on average using the hybrid method, and 176 using the pure threshold method (Metagraph 91 genomes, Bifrost 175 genomes).

**Table 2.**
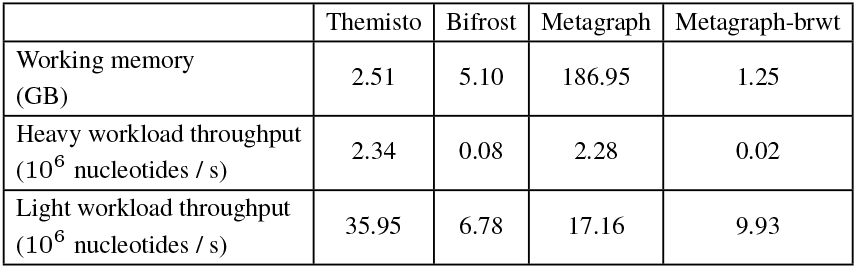
Results of the query benchmark on the largest Salmonella index all three tools were able to construct within 24 hours, consisting of 5596 genomes of S. enterica. The heavy workload consists of reads from isolates of S. enterica. The light workload consists of reads from N. gonorrhoeae. These results are plotted in Figure 8.

|  | Themisto | Bifrost | Metagraph | Metagraph-brwt |
| --- | --- | --- | --- | --- |
| Working memory (GB) | 2.51 | 5.10 | 186.95 | 1.25 |
| Heavy workload throughput ($10^6$ nucleotides / s) | 2.34 | 0.08 | 2.28 | 0.02 |
| Light workload throughput ($10^6$ nucleotides / s) | 35.95 | 6.78 | 17.16 | 9.93 |

The second subplot of Figure 8 shows the query throughput and memory usage of the tools in the low workload experiment. Here, the throughput of all tools increased by at least an order or magnitude. The biggest gain was observed in the binary relation wavelet tree index of Metagraph, which was now two orders of magnitude faster, presumably because the heavy coloring structure is now accessed significantly fewer times than in the heavy workload experiment. However, despite this performance gain for Metagraph, Themisto was still the fastest method by a factor of two.

## 6 Conclusion

We have introduced Themisto a tool backed by highly optimized compressed data structures that allows efficient, scalable indexing of hundreds of thousands of bacterial genomes. Themisto implements a novel hybrid method of pseudoalignment that combines features from existing methods (Bray *et al*., 2016; Holley and Melsted, 2020; Karasikov *et al*., 2020a). Our hybrid method enables queries of both short and long read data by providing adaptable criteria that can be tailored to fit the characteristics of the different sequencing technologies. The method is implemented on top of state-of-the-art indexing techniques which results in construction and query times that are significantly faster than those of existing tools. In our query benchmarks, Themisto always had the highest throughput of all tools tested. Metagraph was almost as fast on the heavy workload, but used 82 times more working memory.

The combination of the color structure and scalability in our new index presents opportunities for developing methods that leverage pseudoalignment not only at the level of individual reference genomes, but also at the level of other (sub)sequences of interest. One particularly interesting avenue for future research is incorporating hierarchical structure in the color set, either in the form of taxonomy, or in the form of gradual progression from genomes to individual genes. In these scenarios, where the number of colors per sequence grows, our results suggest a possibility for developing methods that compress the color sets more effectively.

As a part of larger application pipelines, Themisto has already been instrumental in published metagenomics studies (Tonkin-Hill *et al*., 2022; Mäklin *et al*., 2022) and is incorporated as part of several other ongoing projects, in particular in the study of antimicrobial resistance. As the size and availability of reference genome data sets continues to grow, we expect our improvements to further drive adoption of Themisto as a standard for constructing and querying these large corpora, and the adaptability of our hybrid method to facilitate discovery of entirely new applications for pseudoalignment.

Our indexing results highlight the importance of preprocessing the input data into colored unitigs. Proper preprocessing eliminates redundancy in the data which is irrelevant for *k*-mer based methods such as Themisto. This approach also splits the construction problem into two modular steps which can be worked on independently by separate research groups. The colored unitigs of the input database could also potentially serve as a common data exchange format between different colored de Bruijn graph tools, enabling more efficient interoperability between tools. In this work, we have focused on exact *k*-mer indexing, but we think that recent inexact methods based on the Sequence Bloom Tree (Lemane *et al*., 2022) could potentially be used to approximate Themisto pseudoalignment to a satisfactory precision. It is unclear however how such a method would scale to datasets with a large amount of colors. We are also currently investigating exact methods to simultaneously improve color set size and set intersection time via the use of novel dictionary compression schemes.

## Footnotes

1 These were called *core k-mers* in an earlier version of this manuscript, but were renamed to avoid confusion with the concept of a core genome.

2 See Bussi *et al*. (2021) for a discussion on the choice of *k* in genomics applications.

