## Supplement for "Themisto: a scalable colored *k*-mer index for sensitive pseudoalignment against hundreds of thousands of bacterial genomes"

March 23, 2023

#### 1 Species-colored index of the Blackwell et al. dataset

The constructed index is available at Zenodo at DOI `https://doi.org/10.5281/zenodo.7736981`. The index was built from the 639,981 high-quality genomes from the 661k bacterial genomes dataset of Blackwell et al. [1]. The index contains all distinct 31-mers of the dataset (both strands). There are 71 billion distinct 31-mers in the data (35.5 billion reverse complement pairs). Each k-mer is annotated with the set of species identifiers (colors) that contain that 31-mer. There are 2340 distinct colors in the dataset. The average unitig length in the dataset is 63.5, with a minimum length of 31 and maximum length of 159869.

The de Bruijn graph component of the index (the SBWT data structure) takes 44GB and the colors take 24GB. 99.98% of the color sets were encoded as sparse. The mapping from key $k$-mers to color set identifiers takes 23GB. The color sets themselves take less than a gigabyte. The index was built with the following command:

```
themisto build -i input_file_list.txt --file-colors --reverse-complements -o \
640k_bacteria -m 512000 -t 48 -k 31 --temp-dir temp --verbose -d 20
```

#### 2 Dealing with false positives due to sequence homology with a two-step pseudoalignment pipeline

Since the hybrid method only takes into account $k$-mers that are found in at least one genome in the index, the method may sometimes end up making the pseudoalignment decision on just a small fraction of $k$-mers of the read. This can lead to incorrect pseudoalignments in the following scenario. Suppose we have two species $A$ and $B$ with a significant amount of homology and/or shared mobile genetic elements, and we have built a Themisto index for strains $a_1, \ldots, a_n$ from species $A$. Suppose we have a long read that originates from species $B$, that contains a homologous gene from $A$ and $B$. The hybrid threshold method will likely report this read as compatible to strains of species $A$, even if the sequence surrounding the homologous gene would clearly indicate that the read belongs to species $B$, since those $k$-mers are ignored by the method.

To mitigate this issue, we recommend using a broad index like the one described in Section 1 of this Appendix as the first step in the analysis. This index is constructed to encompass most of the known sequenced bacterial diversity. If the index contains genomes from species $B$ in the previous example, then all the $k$-mers of species $B$ will become relevant in the pseudoalignment, and the erroneous assignment to species $A$ in the example could be avoided. The user could then move to a more specialized index to resolve the strain-level compatibility of each read within the compatible species.

#### 3 Comparison of Kallisto and Themisto pseudoalignment counts

To compare the pseudoalignment of Themisto versus that of Kallisto, we built an index with 100 randomly chosen Salmonella genomes from our Salmonella dataset. Since Kallisto replaces all non-ACGTU nucleotides with random nucleotides, we preprocess the data by deleting all non-ACGTU characters from the genomes. We concatenate all contigs in each genome in order to have Kallisto assign the same color for each contig from the same genome. We used this data to build Themisto and Kallisto index structures. Both tools were run with parameter `-k 31`, and default settings otherwise.

Next, we pseudoaligned the Salmonella isolate sample ERR2693452 against both indexes. We ran themisto with default settings, which means using threshold 1.0 and ignoring unknown $k$-mers. Kallisto was run with settings `--single -l 1 -s 1`. With these settings, the pseudoalignment algorithm of Themisto should in theory match that of Kallisto, with the exception of the unitig skipping heuristic employed in Kallisto.

To analyze the results, we collected the number of hits to each color from both tools. For each color, we computed the hit ratio, i.e. the number of reads compatible with that color according to Kallisto divided by the number of reads compatible with that color according to Themisto. The distribution of these ratios over all colors is plotted in Figure 1. We see that the unitig skipping heuristic of Kallisto has a false positive rate of around 0.03% on this dataset, assuming there are no other implementation differences between the tools.

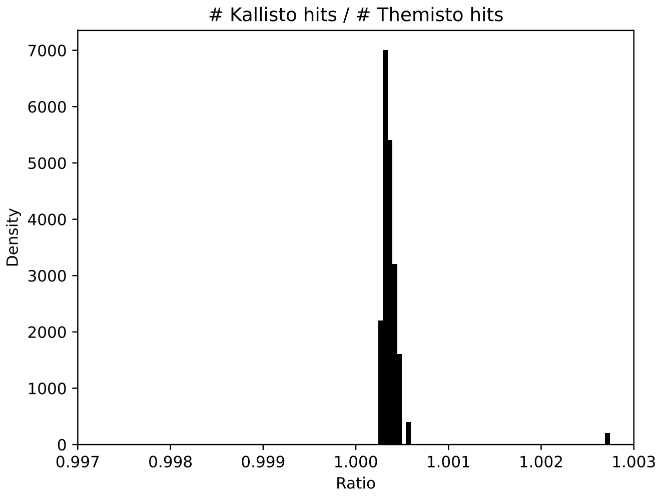


Figure 1: Ratio of pseudoalignment counts between Kallisto and Themisto.

### 4 Salmonella experiment command line parameters

Here we list the command line parameters used to construct and query the Salmonella index structures in the main paper.

#### 4.1 Bifrost parameters

The construction was run with parameters `build -r -c -t 48 -k 31`. The queries were ran with `-t 48 --ratio-kmers 0.8`.

#### 4.2 Metagraph parameters

##### 4.2.1 $k$-mer index construction

The $k$-mer index was build using the parameters `build -k 31 --mem-cap-gb 1000 -p 48`.

##### 4.2.2 Uncompressed row-major color representation

The uncompressed row-major representation was constructed using the command line parameters `annotate -v --anno-filename -p 48 --anno-type row --mem-cap-gb 1000 --disk-swap`. The queries were run with `--discovery-fraction 0.8 -p 48`.

##### 4.2.3 RowDiff<Multi-BRWT> color representation

Following the user manual, we constructed the RowDiff<Multi-BRWT> representation via the column-major representation with the following sequence of subcommands:

1. `annotate -v --anno-filename -p 48 --mem-cap-gb 1000`
2. `transform_anno -v -p 48 --anno-type row_diff --row-diff-stage 0 --mem-cap-gb 1000`
3. `transform_anno -v -p 48 --anno-type row_diff --row-diff-stage 1 --mem-cap-gb 1000`
4. `transform_anno -v -p 48 --anno-type row_diff --row-diff-stage 2 --mem-cap-gb 1000`
5. `transform_anno -v -p 48 --anno-type row_diff_brwt --mem-cap-gb 1000 --greedy`
6. `relax_brwt -v -p 48 --relax-arity 32`

Note that unlike in the regular row-major construction, the commands were run without `--disk swap` because that command caused an error during construction in this pipeline. The queries were run with `--discovery-fraction 0.8 -p 48`.

#### 4.3 Themisto parameters

Themisto was constructed with `build -k 31 --temp-dir temp -m 1000000 -t 48 -v`. The queries were run with `--threshold 0.8 --ignore-unknown-kmers -t 48 --buffer-size-megas 0.1`.

### 5 Complete data from the index construction benchmark.

This table lists the construction running time, peak memory, and the size of the de Bruijn graph (DBG) and color components of the index structures on disk. Missing data for runs which exceeded 24 hours are marked with NA.

| | Themisto | | | | Bifrost | | | | Metagraph | | | | Metagraph-brwt | | | |
|---|---|---|---|---|---|---|---|---|---|---|---|---|---|---|---|---|
| Number of genomes | Time (h) | Mem (GB) | DBG (GB) | Colors (GB) | Time (h) | Mem (GB) | DBG (GB) | Colors (GB) | Time (h) | Mem (GB) | DBG (GB) | Colors (GB) | Time (h) | Mem (GB) | DBG (GB) | Colors (GB) |
| 1 | 0.01 | 1.42 | 0.01 | 0.07 | 0.00 | 0.11 | 0.00 | 0.00 | 0.00 | 0.38 | 0.02 | 0.00 | 0.00 | 0.38 | 0.02 | 0.00 |
| 2 | 0.01 | 1.50 | 0.01 | 0.00 | 0.00 | 0.12 | 0.00 | 0.00 | 0.00 | 0.46 | 0.02 | 0.00 | 0.00 | 0.47 | 0.02 | 0.00 |
| 5 | 0.01 | 1.61 | 0.01 | 0.01 | 0.00 | 0.17 | 0.01 | 0.00 | 0.00 | 0.77 | 0.02 | 0.01 | 0.00 | 0.56 | 0.02 | 0.00 |
| 10 | 0.02 | 1.88 | 0.01 | 0.03 | 0.00 | 0.20 | 0.01 | 0.01 | 0.01 | 1.16 | 0.02 | 0.03 | 0.01 | 0.72 | 0.02 | 0.00 |
| 21 | 0.02 | 2.17 | 0.02 | 0.06 | 0.01 | 0.25 | 0.02 | 0.01 | 0.01 | 1.81 | 0.03 | 0.07 | 0.01 | 1.10 | 0.03 | 0.00 |
| 43 | 0.03 | 2.51 | 0.02 | 0.07 | 0.01 | 0.36 | 0.02 | 0.02 | 0.02 | 2.73 | 0.03 | 0.16 | 0.03 | 2.07 | 0.03 | 0.00 |
| 87 | 0.04 | 3.02 | 0.03 | 0.11 | 0.02 | 0.43 | 0.03 | 0.04 | 0.05 | 4.43 | 0.04 | 0.37 | 0.07 | 3.78 | 0.04 | 0.01 |
| 174 | 0.06 | 3.73 | 0.04 | 0.16 | 0.05 | 0.68 | 0.04 | 0.08 | 0.09 | 7.74 | 0.04 | 0.82 | 0.13 | 7.10 | 0.04 | 0.02 |
| 349 | 0.08 | 4.18 | 0.05 | 0.23 | 0.10 | 0.90 | 0.06 | 0.17 | 0.19 | 14.04 | 0.05 | 1.82 | 0.27 | 13.67 | 0.05 | 0.04 |
| 699 | 0.12 | 5.44 | 0.07 | 0.39 | 0.23 | 1.38 | 0.09 | 0.38 | 0.51 | 26.74 | 0.07 | 4.00 | 0.72 | 26.37 | 0.07 | 0.10 |
| 1398 | 0.17 | 9.06 | 0.09 | 0.64 | 0.47 | 2.40 | 0.12 | 0.65 | 1.34 | 51.53 | 0.09 | 8.72 | 1.60 | 51.53 | 0.09 | 0.21 |
| 2796 | 0.27 | 15.99 | 0.12 | 1.19 | 1.17 | 4.25 | 0.17 | 1.58 | 3.62 | 101.51 | 0.11 | 18.90 | 4.04 | 101.51 | 0.11 | 0.48 |
| 5593 | 0.40 | 30.08 | 0.15 | 2.29 | 2.76 | 8.16 | 0.22 | 3.00 | 9.98 | 201.55 | 0.13 | 40.80 | 9.11 | 201.55 | 0.13 | 1.04 |
| 11186 | 0.69 | 56.68 | 0.22 | 5.03 | 6.71 | 13.24 | 0.32 | 4.49 | NA | NA | NA | NA | 19.96 | 401.58 | 0.17 | 2.40 |
| 22373 | 1.25 | 107.39 | 0.28 | 11.12 | 15.20 | 26.78 | 0.43 | 11.59 | NA | NA | NA | NA | NA | NA | NA | NA |
| 44746 | 2.31 | 168.20 | 0.37 | 25.59 | NA | NA | NA | NA | NA | NA | NA | NA | NA | NA | NA | NA |
| 89492 | 4.61 | 291.55 | 0.51 | 59.83 | NA | NA | NA | NA | NA | NA | NA | NA | NA | NA | NA | NA |
| 178984 | 9.37 | 542.08 | 0.70 | 140.42 | NA | NA | NA | NA | NA | NA | NA | NA | NA | NA | NA | NA |
